# Enhanced tameness by *Limosilactobacillus reuteri* from gut microbiota of selectively bred mice

**DOI:** 10.1101/2024.03.11.584526

**Authors:** Bhim B. Biswa, Hiroshi Mori, Atsushi Toyoda, Ken Kurokawa, Tsuyoshi Koide

## Abstract

Domestication alters animal behaviour, primarily their tameness. In this study, we examine the effect of gut bacteria on mouse tameness. We previously conducted selective breeding for active tameness, defined as the motivation to approach a human hand, using genetically heterogeneous mice derived from eight wild inbred strains. We examined gut microbiota in the selectively bred mice by analysing faecal samples from 80 mice through shotgun metagenomic analysis. In the current study, we found that the selectively bred mice exhibit higher levels of active tameness as well as higher levels of blood oxytocin, which plays a key role in social behaviours. Selection for tameness did not substantially alter the taxonomic or functional diversity of the gut microbiota. However, we observed an increased abundance of *Limosilactobacillus reuteri* in the selected groups and higher pyruvate levels in their plasma. We isolated *L. reuteri* strains secreting extracellular pyruvate from mice faeces and administrated the cultured bacteria through drinking water. Mice treated with *L. reuteri* showed higher colonization of the bacteria in the gut, as well as higher levels of active tameness behaviour and blood oxytocin. Additionally, we generated 374 high-quality metagenome-assembled genomes (MAGs) of bacteria across 11 phyla. This collection includes 27 novel species level bacterial MAGs not previously known to exist in the mouse gut. This study elucidates the potential role of *L. reuteri* in the animal domestication process and explores the underlying mechanisms that may influence this process.

## Introduction

Domestication is a sustained multigenerational process by which captive animals adapt to humans and their environment. Several factors have been proposed to contribute to the adaptation of animals to captive environments. Most adaptation processes are explained by genetic changes; however, other factors such as certain experiences and environments, proximity and handling by humans^1–3^, temperature^4–6^, and food^7^ may also contribute to the adaptation process. Therefore, domestication is a combination of genetic changes that occur over generations and of environmentally induced ontogenic events^8^. Multiple studies have shown that gut microbiota influence host behaviour, brain development, and cognition. The metabolites produced by the microbiota alter the host’s metabolic pathways or influence their immune system^9,10^. Several studies have explored the association between gut microbiota and domestication in animals, such as mice^11^, horses^12,13^, buffalo^14^, turkeys^15^, chicken^16^, and several animal species together^17^. These studies have identified multiple species of bacteria that may influence the process of animal domestication. However, such studies are limited by the risk of contamination of the gut microbiota due to a non-sterile housing environment, the lack of common ancestors of test and control animals, differences in feed and water between test animals, crossover contamination from other animal species, and the use of antibiotics during rearing.

These challenges can be mitigated by using animals bred in a controlled setting with a singular focus on phenotypic selection and well-documented lineages. The WHS mice, which have been selectively bred for tameness are a good example of this approach. Tameness is a behavioural phenotype necessary for animals to adapt to humans and presents in two types: the motivation to approach humans (active tameness) and a reluctance to avoid humans (passive tameness). To increase active tameness, mice that exhibited a relatively high motivation to approach humans in the active tameness test, which evaluates levels of active tameness in mice^18^, were bred selectively using wild-derived heterogeneous stock (WHS) mice. A founder group of WHS mice was developed by crossing eight different wild inbred strains originating from different countries and from three subspecies groups of *Mus musculus domesticus, castaneus, and musculus*^19,20^. A group of WHS was bred with 16 pairs of mice in each generation to maintain high genetic diversity. Four groups of WHS were generated from the common founder WHS group by splitting them into two selected (S1 and S2) and two non-selected WHS groups (C1 and C2) (Fig. 1A). Possible genetic components of active tameness have been reported in these WHS mice^20,21^, suggesting that genetic factors play a major role in the control of tameness; however, the role of the gut microbiota still needs to be investigated. All WHS mice were kept in the same animal room but groups were kept isolated in different breeding racks. Furthermore, all the mice including the founder wild inbred strains were kept in specific pathogen-free (SPF) conditions and were given radiation-sterilised food and ultrafiltration-sterilised water. Therefore, any gut microbiota difference between selected and non-selected groups can be directly associated with the selection pressure on the host.

**Fig. 1.**
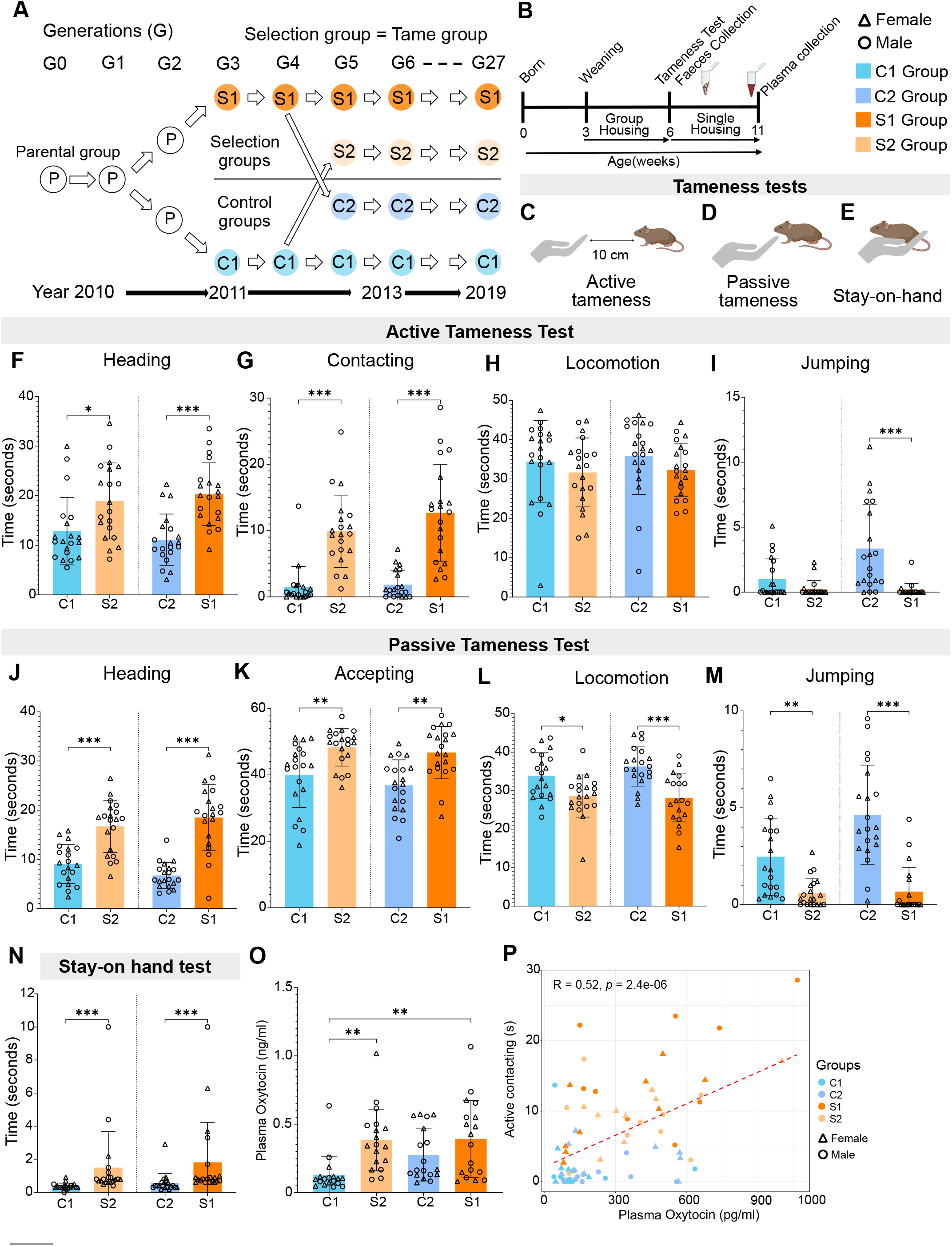
Tameness and plasma oxytocin concentration is higher in selected mice compared to control. (A) Scheme of wild heterogenous stock generation; (B) Experimental timeline, (C-E) tameness test illustrative depiction, (C) active tameness test, (D) Passive tameness test, (E) stay- on-hand test, (F-I) active tameness test results, (F) heading, (G) contacting, (H) locomotion, (I) jumping, (J-M) passive tameness test results, (J) heading, (K) accepting, (L) locomotion, (M) jumping, (N) stay on hand test, (O) Plasma oxytocin concentration (P) Pearson correlation between oxytocin concentration and active contacting time. N for tameness test = 80 (20 in each group with 10 male and 10 female); N for oxytocin ELISA = 72 (18 in each group with 9 male and 9 female). (*p<0.05; **p<0.01; ***p<0.001). Bar graphs show means ± SD with individual data points.

In our study, to characterize the gut microbiota in WHS mice, we conducted metagenomic analysis using genomic DNAs obtained from the faeces of animals from the selected and non- selected groups. An intervention strategy was then used to introduce specific bacteria into the drinking water of the mice to examine changes in their tameness behaviour.

## Materials and Methods

### Experimental conditions and mice

All experiments in this study were conducted in strict accordance with the guidelines and protocols sanctioned by the “Committee for Animal Care and Use” at the National Institute of Genetics (permit numbers: R4-25 and R5-7). The mice used in our experiments were bred and maintained in specific pathogen-free (SPF) conditions within the animal facility at NIG. All equipment used for animal maintenance, including standard-sized transparent polycarbonate cages, cage tops, paper bedding materials, and other essential supplies, underwent sterilization through autoclaving. The mice were provided with a standard chow diet, CE-2 (CLEA Japan, Inc., Tokyo, Japan), which was radiation-sterilized at 15 KGy. Both food and ultrafiltration sterilized water (supplemented with 3ppm chlorine) were availed to the mice ad libitum. The mice were housed in a temperature-controlled room at 23±2°C with a humidity range of 50±10% and a 12-h light-dark cycle with lights switched on at 6:00 am. Mice were weaned at approximately 3–4 weeks of age and were co-housed with their same-sex littermates until the tameness test was conducted. During cage changes and behavioural tests, each mouse was gently handled by its tail using tweezers covered with a silicone tube to minimize stress. All personnel involved in the experiments wore polyester suits, non-woven fabric masks, and double gloves (cloth gloves under latex gloves) to maintain a sterile environment. In all experiments, we took careful measures to prevent microbiome cross-contamination. Cages housing mice from different groups were not placed in close proximity to each other. This precaution ensured that the microbial environments of each group remained distinct and uncontaminated by other groups. The first set of experiments was carried out using mice from the 27^th^ generation of the WHS. The mouse stocks used in this study, S1, S2, C1, and C2, are available from the following website https://grc.nig.ac.jp/mgrlab/index.xhtml.

### Selective breeding of WHS

WHS mice were generated by crossing eight wild strains, BFM/2Ms, PGN2/Ms, HMI/Ms, NJL/Ms, BLG2/Ms, KJR/Ms, CHD/Ms, and MSM/Ms, derived from different countries to expand the genetic heterogeneity. The founder stock, which is kept with 16 pairs of breeding, was split into two groups in the third generation to create S1 and C1, and, henceforth, S1 was selectively bred for higher active tameness. In the fifth generation, another selected group, S2, was split from C1, and another non-selected group, C2, was split from S1. Thereafter, S2 was selectively bred for a higher level of active tameness and C2 was maintained as the non-selected group. Mice used for mating were selected from each of the five females and five males offspring of each pair based on the highest contacting score in the active tameness test, and if two or more animals showed the same highest contacting score, the individual with the higher heading score in the same test was selected.

### Tameness test

The tameness test was performed at six weeks of age, as described in a previous study^18^ during the light period between 14:00-17:00. The test was conducted in an open-field arena (40×40×40cm) made of grey polyvinyl chloride (O’Hara & Co. Ltd., Tokyo, Japan) with the floor illuminated with approximately 100 lux at the centre^22,23^. The tameness test was a three-min protocol divided into one minute each of the active and passive tameness tests and three trials of the stay-on-hand test. In the active tameness test, the experimenter’s hand was kept 10cm away from the test mouse for 1 min with the fingers moving slightly and rhythmically to observe the animal’s motivation to approach and contact the human hand. In the passive tameness test, the experimenter attempted to touch the mouse with their hand, and mouse behaviour was recorded for 1 min. This test examined how long the mice could tolerate being touched by a human hand. In the stay-on-hand test, each mouse was picked from its tail using silicon-coated tweezers, placed on the experimenter’s hand, and stroked gently until it attempted to get off the hand. All tests were video-recorded and analysed by another experienced researcher, M. Nihei, using tanaMove (zenodo.org/records/10829003)^24^ and scores were assigned to nine different parameters. Video analysis of all tameness tests was blinded with respect to the grouping of mice. The detailed step-by-step process has been outlined in previous studies^22,23^.

### Tameness test parameter clustering

The analysis was performed in R Ver.4.3.2. The tameness test results for all 80 samples were normalized using “min-max” normalization. Clustering was executed based on the correlation distance, employing the Ward method via the “hclust” function. The statistical significance of the identified clusters was assessed using the “pvclust” function from the R package pvclust (Ver.2.2.0)^25^, setting the significance level at 5% for each clustering analysis.

### Mice faeces collection

In all experiments, mouse faecal samples were collected following the tameness test, ensuring consistency in timing between 17:00 and 18:00 on the same day. To maintain sterility and minimize environmental variables, the test mice were housed in autoclaved, sterile cages devoid of any bedding material. This setup allowed for natural defecation without external prompting. The faeces were carefully collected in an Eppendorf tube using sterile aluminium foil and immediately stored at −80°C.

### Blood serum and plasma collection

Blood plasma was obtained when mice reached 11 weeks of age. For the collection procedure, the mice were first anaesthetized using an intraperitoneal (IP) injection of pentobarbital at a dosage of 50mg/kg body weight (Tokyo Chemical Industry Co. Ltd., Japan). Following anaesthesia, blood samples were drawn via cardiac puncture using a syringe pre-treated with heparin to prevent coagulation. The collected blood was then transferred into heparinized Eppendorf tubes. These tubes were allowed to stand undisturbed at room temperature for 30min to enable the separation of blood components. Subsequently, the tubes were centrifuged at 10,000G for 10min at a temperature of 4°C. For serum isolation, the same protocol was used, excluding heparin treatment.

### Metagenomic DNA isolation from mice faeces

In our modified approach to metagenomic DNA isolation, we utilized protocol #6 from Costea et al.^26^, incorporating the QIAamp Fast DNA Stool Mini Kit (QIAGEN GmbH, Hilden, Germany) for DNA extraction. The process began with treating 180–220mg of faecal samples with 1ml of lysis buffer (500mM NaCl, 50mM Tris-HCl at pH 8.0, 50mM EDTA, and 4% sodium dodecyl sulfate). This mixture was homogenized for 5min using a handheld homogenizer (Leda Trading Corp.), followed by the addition of 10µl proteinase K and sterile zirconia beads (1.0mm, 20-30; BioSpec, Inc., USA). For thorough mixing, the sample was vortexed at maximum speed for 10 min using a Vortex-Genie 2 mixer (MO BIO Laboratories, Inc., USA) and then incubated at 95°C for 15min. Post incubation, the sample was centrifuged at 16,000xg for 5min at 4°C, and the supernatant was transferred to a new 2mL tube. The remaining pellet was subjected to a second round of lysis, resuspended in 300µL of the lysis buffer, and processed as previously described. The combined supernatants from both lysis steps were then treated with 260µL of 10M ammonium acetate, followed by vortexing and incubation on ice for 5min. The solution was centrifuged at 16,000xg for 10min at 4°C, and the resulting supernatant was equally divided into two 1.5 mL tubes, each containing 750 µL isopropanol. These tubes were then incubated on ice for 30 min and centrifuged at 16,000 x g for 15 min. The supernatant was discarded, and the pellet was washed with 0.5 mL of 70% ethanol and left to air-dry. Finally, the dried pellets were reconstituted in 100 µL of TE (Tris-EDTA) buffer, and the aliquots were pooled. We added 2 μL DNase-free RNase (10 mg/mL) to the pooled DNA solution and incubated it at 37°C for 15 min. Then, 10 μL proteinase K and 200 μL buffer AL were added, vortexed for 15 s, and incubated at 70°C for 10 min. Subsequently, 200 μL of 96–100% ethanol was added to the lysate, vortexed, and passed through a QIAamp spin column, and centrifuged for 1 min. The filtrate was discarded and the column was sequentially washed with 500 μL each of Buffer AW1 and AW2. Following a final centrifugation, 100 μL AE Buffer was used to elute DNA into a new 1.5 mL tube. A spectrophotometer (NanoVue Plus) was used for the quality check. A Qubit 2.0 Fluorometer was used for precise quantification following the manufacturer’s instructions. DNA integrity was checked using 0.8% agarose gel electrophoresis. The DNA was then stored at −20°C for future use.

### Plasma metabolites analysis

For metabolomic analysis, samples were analysed by Human Metabolome Technologies, Inc. (https://humanmetabolome.com/jpn/). A basic scan was performed to detect more than 1000 water-soluble metabolites. The protocol for this analysis has been described previously^27–30^. Briefly, metabolome analysis (CE-TOFMS) was performed on 12 randomly selected mouse plasma samples in two modes for cationic and anionic metabolites.

### Limosilactobacillus reuteri isolation

Cecum and faecal samples were collected from mice of selected group (S1), mixed, and then serially diluted. This diluted mixture was cultured on *L. reuteri* Isolation Media (LRIM)^31^ agar plates, with raffinose (Tokyo Chemical Industry Co. Ltd, Japan) as the sole carbon source. The plates were incubated for 48h at 45°C under anaerobic conditions (5% CO2). To maintain the anaerobic condition, plates were stored in a sealed box containing one AnaeroPack sachet (Mitsubishi gas chemical company, Inc, Japan). For the isolation of pure cultures, selected colonies were first streak plated twice on LRIM plates. This was followed by third streak plating on De Man–Rogosa–Sharpe (MRS) (Merck-Millipore, USA) agar plate.

### Limosilactobacillus reuteri phylogenetic analysis

All isolated colonies were cultured in MRS media, after which DNA was extracted using the NucleoSpin Microbial DNA kit (Takara Bio Inc., Shiga, Japan), according to the manufacturer’s protocol. To amplify the full 1.6 kbp region of the 16S rDNA, polymerase chain reaction (PCR) was performed using the universal primers bak4 (5’-AGGAGGTGATCCARCCGCA-3’) and bak11w (5’-AGTTTGATCMTGGCTCAG-3’)^32,33^. Cycling conditions were 95°C for 5min, followed by 35 cycles of 95°C for 15s, 60°C for 30s and 72°C for 2min, and final extension for 7min at 72°C. The PCR cycling conditions included an initial denaturation at 95°C for 5 min, followed by 35 cycles of 95°C for 15s, 60°C for 30s, and 72°C for 2min, with a final extension at 72°C for 7min. The PCR products were then purified and subjected to Sanger sequencing.

Taxonomic identification of obtained sequences was done using NCBI BLASTn^34^. To infer phylogeny, along with our sequences, we first obtained 16S rRNA gene sequence of multiple *Limosilactobacillus* species from NCBI along *Lactobacillus helveticus* as outgroup. All sequences were aligned using the MAFFT online server with “FFT-NS-I” command^35,36^. Using Model finder^37^, we found “TPM3u+F+I” as the best-fit model for out data. We used IQ-TREE 2^38^ with 1000 bootstrap to construct a maximum likelihood tree. This tree was edited using iTOL online server^39^.

### Pyruvate, L-Lactate, and D-Lactate secretion assay

In this experiment, *L. helveticus* (JCM1120), sourced from the Japan Collection of Microorganisms (JCM), served as the positive control for pyruvate production. All bacterial strains, including the control, were cultured in Gifu Anaerobic Broth (GAM) medium (Nissui Pharmaceutical, Japan), which was supplemented with 1% glucose. The incubation was conducted for 24 h at 37°C under anaerobic conditions. To measure the concentrations of pyruvate, L-Lactate, and D-Lactate in the media, we utilized biochemical assay kits from Cayman Chemicals (Michigan, USA), strictly adhering to the manufacturer’s protocol. All three assays were conducted in triplicates (technical replication).

### Limosilactobacillus reuteri administration

All bacterial strains were cultured in MRS media for 24 h. Following this cultivation, the optical density (OD) at 590nm was measured to estimate the number of bacteria in the solution, using a standard curve. The bacteria were then washed with PBS, resuspended in PBS, and stored at −80°C until use. We cultured a fresh batch of bacteria every week to maintain viability. PBS (vehicle) or bacteria (∼1 x 10^8^ CFU/Cage/day) was added to the drinking water of the mice each day between 17:00 and 18:00. Mice from the 37^th^ generation of C1 group were used. For the drinking water administration and measurements, 15-mL glass test tubes stoppered with silicone caps with double ball-bearing nozzles, Drinko-measurer DM-G1 (O’hara & Co. Ltd., Tokyo, Japan), were used daily. The administration of *L. reuteri* began immediately post-weaning, at approximately 3 weeks of age, and was continued for 21 days. To minimize isolation stress, mice were housed in same-sex pairs. Faecal samples were collected from each individual both before the start and at the end of the experimental period. In addition, we monitored and recorded the daily water consumption of each cage and measured the body weight of the mice at the end of the experiment. After the 21- day bacterial administration period, a tameness test was conducted. This was followed by the collection of faecal and plasma samples for further analysis.

### qPCR quantification of Limosilactobacillus reuteri and Lactobacillus helviticus

Quantitative PCR (qPCR) was conducted using a Thermal Cycler Dice Real Time System III (Takara Bio Inc., Shiga, Japan). For each 25μL reaction, TB Green Premix Ex Taq II (Tli RNaseH Plus) (Takara Bio Inc., Shiga, Japan) and gene-specific primers at a concentration of 1 μM were used. The reaction mixtures included 5ng of metagenomic DNA, the quantity of which was determined using Qubit. The cycling conditions were set at 95°C for 5min, followed by 40 cycles of denaturation at 95°C for 20s, annealing at 62°C for 20s, and extension at 72°C for 30s. After the PCR cycles, a melt curve analysis was performed in the range of 60–95°C using the default settings of the system. The primers used in the qPCR were as follows: Universal bacterial primers EUB338 (5’-ACTCCTACGGGAGGCAGCAG-3’)^40^, and EUB518 (5’- ATTACCGCGGCTGCTGG-3’)^41^; *L. reuteri* -specific 16S-23S rRNA gene spacer primers, sg- Lreu-F (5’-GAAGATCAGTCGCAYTGGCCCAA-3’), and sg-Lreu-R (5’- TCCATTGTGGCCGATCAG-3’)^42^; *L. helveticus* hsp60 specific primers, F1LHelHsp (5’- CTTTGATCGCTGATGCTATGGAAAAGGTTGGTC-3’), and R1LHelHsp (5’- GATCAACAATGACTTGCCTTGTTGAACAATTTC-3’)^43^. The qPCR analysis was based on the ΔCt of the specific primers normalized to the ΔCt of the universal primers (ΔΔCt method). The qPCR analysis was performed in triplicates (technical replication).

### Mice serum pyruvate and oxytocin concertation

To measure the concentrations of pyruvate in mice serum, we utilized biochemical assay kits from Cayman Chemicals (Michigan, USA), strictly adhering to the manufacturer’s protocol. Due to the limited number of sample wells, 10 randomly selected samples (5 males and 5 females) were analysed. To measure oxytocin concentration, we used Oxytocin ELISA kit (Enzo Life Sciences, USA), following manufacturer’s instructions. For baseline oxytocin in WHS mice, the ELISA plate’s well constraints meant we analysed 18 plasma samples (9 male and 9 female) from each group, totalling 72 samples. For looking at the effect of *L. reuteri* treatment on blood oxytocin levels, *L. helveticus*-treated samples were excluded. Both assays were performed in duplicates (technical replication).

### Shotgun metagenomic sequencing

Library preparation and sequencing were performed at the Advanced Genomics Center of the National Institute of Genetics. Briefly, genomic DNA was fragmented to an average size of 500bp using a DNA shearing system (M220 focused ultrasonicator; Covaris Inc., MA. USA). A paired-end library was constructed using a TruSeq DNA PCR-Free Library Prep kit (Illumina, CA, USA) and size-selected on an agarose gel using a Zymoclean Large Fragment DNA Recovery Kit (Zymo Research, CA. USA). The final library was subjected to paired-end sequencing using an Illumina NovaSeq 6000 sequencer with a read length of 150bp.

### Bioinformatics analysis of metagenomic sequence data

#### Quality control of metagenomic sequences

All sequencing reads obtained were quality-filtered using KneadData(Ver.0.10.0) (https://github.com/biobakery/kneaddata), where low-quality sequences and any contamination from PhiX, host (mouse_C57BL_6NJ.1), human genome (hg37dec_v0.1.1), or transcriptome (human_hg38_refMrna.1) were discarded. KneadData uses Trimmomatic(Ver.0.33)^44^ to remove adapters, TRF^45^ to remove repetitive sequences, and Bowtie2(Ver.2.4.5)^46^ to remove contaminants.

#### Taxonomic analysis

Taxonomic analysis of the gut microbiome was conducted using Kraken 2 (Ver.2.1.2)^47^. We employed the full NCBI/RefSeq database made by Wright et al. (NCBI RefSeq Complete V205 (1189 GB))^48^ to ensure comprehensive species identification within the gut samples. Default parameters were used for Kraken 2, with a confidence score threshold set at 0.15. This threshold was chosen to strike a balance between the robustness of species identification and maximizing the percentage of sequences utilized in the analysis. Further refinement of the Kraken 2 output was achieved through Bayesian re-estimation using Braken2(Ver.2.6.2), also utilizing its default settings^49^.

#### Functional Analysis

For functional analysis in our study, we utilized HUMAnN3(Ver.3.0.1)^50^ employing the ‘–bypass- prescreen’ option. We concatenated the filtered forward and reverse sequences from each sample and mapped them to the comprehensive ChocoPhlAn database(Ver.201901b). Sequences that remained unassigned after this step were further analysed against the UniRef90 database. To identify specific metabolic pathways, we employed the MetaCyc database^51^. We obtained Copies per Million data for each gene family and pathway using the ’humann_renorm_table’ command.

#### Abundance and Diversity Analysis

Diversity analysis and graph generation in our study were carried out using the MicroEco package^52^ We normalized the abundance data obtained from Braken2 by rarefying it to the size of the smallest sample^53^. To assess the statistical significance of alpha diversity, we employed the Kruskal–Wallis Rank Sum test. Beta diversity was analysed based on Bray-Curtis dissimilarity; analysis was complemented by principal coordinate analysis, which was applied to both taxonomic and functional data. In our correlation analysis, we first conducted a random forest classification using ‘randomForest’ package^54^, utilizing mouse group information (C1, C2, S1, and S2) on the normalized Bracken2 bacterial abundance data, adjusting the p-value using the Benjamini-Hochberg False Discovery Rate (FDR) method. Afterward, we normalized the scores of tameness parameters through min-max normalization. This normalized tameness scores were then subjected to Pearson correlation with the top 40 significant taxa identified in the random forest analysis using ‘pheatmap’ package^55^ .

#### MaAsLin2 association analysis

For association between taxonomic abundance with each mice group, we use Bracken2 abundance data. Initially it was normalized using ’Total sum scaling’, then transformed into a log scale, after which association analysis was performed. In association analysis between functional pathways, we did not perform additional normalisation or transformation as HUMAnN3’s output was already normalized. In both analyses, we used a linear model and compared S1 and S2 groups to their respective controls.

#### MAG generation

Default settings were used in different software, unless otherwise specified. Filtered sequences were assembled using metaSPAdes(k=21,33,55)(Ver.3.15.4)^56^ for a single assembly or MEGAHIT(--k-min 21 --k-max 77)(Ver.1.2.9)^57^ for co-assembly. Multiple binners with different binning strategies were used. The MEGAHIT assembly was binned using MetaCoAG(Ver.1.0)^58^, MaxBin2(Ver.2.2.7)^59^, and MetaBAT2(Ver.2.15)^60^. This was followed by the generation of a nonredundant set of bins using DAStool(search_engine diamond)(Ver.1.1.4)^61^. For the metaSPAdes-generated assembly, we utilized SemiBin(multi_easy_bin)(Ver.0.7.0)^62^ and VAMB(multisplit)(Ver.3.0.8)^63^. All bins produced were subjected to contamination filtering using MDMcleaner(Ver.0.8.3)^64^. Bins devoid of chimeric sequences, as confirmed by GUNC(Ver.1.0.5)^65^, and those with MAG quality scores above 0.5 were retained for further analysis. These filtered bins were then evaluated for contamination and completeness using CheckM(lineage_wf)(Ver.1.2.0)^66^. To create a nonredundant bin set, we employed the “dereplicate” command of dRep(Ver.3.0.0)(-comp 50, -con 10, -pa 0.90, -sa 0.95, -nc 0.30, and -cm larger)^67^. Finally, the taxonomic classification of all bacteria and the generation of the species tree were performed using GTDB-Tk(classify_wf)(Ver.2.1.1) with the “r207_v2” database^68^. MAGs phylogenetic tree was generated using FastTree (Ver.2.1.11)^69^ within GTDB-Tk using the “infer” command. The generated tree was then refined and edited using the Interactive Tree of Life(iTOL) tool^39^.

#### MAG analysis

To identify novel MAGs, we utilized the ’compare’ command of dRep(Ver.3.0.0)(-pa 0.90 -sa 0.95 -nc 0.30 -cm larger)^67^. This analysis was conducted on a combined database that included the 1573 mouse gut MAGs previously compiled by Kieser et al.^70^ along with the 374 MAGs generated in our study. We classified our MAGs as novel if they did not cluster with any of the existing MAGs in this integrated database at 95% ANI, thereby distinguishing them as unique at species level. To quantify the relative abundance of each MAG, we employed CoverM(Ver.0.6.1) (https://github.com/wwood/CoverM) and used minimap^71^ to map reads to our MAG database. All 27 novel MAGs were annotated using DFAST^72^.

## Statistical analysis

GraphPad Prism (Ver.10.1.2) was used for all statistical analyses. To evaluate the normality of the dataset, the Shapiro–Wilk test was utilized. For data that followed a normal distribution, one-way ANOVA was applied, with subsequent Tukey’s test for multiple comparisons. In cases where data did not conform to normal distribution, the Kruskal–Wallis test was employed, followed by Dunn’s test for multiple comparisons. To check the effect of sex on tameness behaviour, a two-way ANOVA was performed followed by Tukey’s test for multiple comparisons. For statistical association analysis of plasma metabolites, taxonomic abundance, and functional abundance, MaAsLin2(Ver.1.12.0)^73^ was used with a Benjamini-Hochberg q-value of 0.25 following previous microbiome studies^74,75^. For beta-diversity, Wilcoxon rank-sum test, two-sided was performed. For MAGs abundance, two tailed Mann–Whitney U test was performed. For daily water intake, serum pyruvate and oxytocin concentration, a two-way ANOVA was performed followed by Tukey’s test for multiple comparisons. For mice body weight, the Kruskal–Wallis test was employed, followed by Dunn’s test for multiple comparisons.

## Results

### Selective breeding leads to increase in active tameness behaviour and blood oxytocin levels

At the 27^th^ generation of breeding, we examined tameness-related behaviours in the selected and non-selected groups of mice (Fig.1A). We conducted tests of active tameness, passive tameness, and stay-on-hand tests on 80 mice: 10 male and 10 female mice for each group (Fig.1B-E). As S1 and C2, and S2 and C1 are genetically closely related (Fig.1A), we compared the data between the two pairs to determine the effect of selective breeding on active tameness compared to the non-selected control, hereafter. Data on nine behavioural parameters (Table S1)—heading, contacting, locomotion, and jumping in the active tameness test (Fig.1F-I); heading, accepting, locomotion, and jumping in the passive tameness test (Fig.1J-M); and median staying time in the stay-on-hand test (Fig.1N)—were obtained. Given that there was no effect of sex in the two-way analysis of variance (Two-way ANOVA; Table S2), data from both sexes were combined in all tameness analyses. From the clustering analysis of all nine parameters, we identified two significant clusters that categorize the parameters as either positively or negatively associated with tameness (Fig.S1). Regarding active contacting, a parameter of selective breeding, both S1(p<0.001) and S2(p<0.001) exhibited significantly longer contact durations compared to their respective controls (Fig.1G; Movies S1, S2). Other tameness-related parameters, such as heading in the active tameness test (Fig.1F)(S1,p<0.001; S2,p<0.05), heading (Fig.1J)(S1,p<0.001; S2,p<0.001) and accepting (Fig.1K)(S1,p<0.001; S2,p<0.01) in the passive tameness test were higher in the selected groups than in the non-selected groups. In the stay-on-hand test, mice of the selected groups stayed on the hand longer than controls (Fig.1N)(S1,p<0.001; S2,p<0.001). In contrast, parameters that are characteristic of wild mice, such as active jumping (Fig.1I)(C2,p<0.001), passive locomotion (Fig.1L)(C2,p<0.001; C1,p<0.05), and passive jumping (Fig.1M)(C2,p<0.001; C1,p<0.01), were higher in the non-selected groups than in the selected groups. Collectively, S1 and S2 showed higher active and passive tameness than the non-selected group.

Previous studies on WHS mice showed that selected groups exhibit higher level of social behaviours as well as higher expression of hippocampal *Oxtr*, a receptor gene for oxytocin which play crucial role in social bonding^76^ in these mice^20,77^. To see how selective breeding on tameness affect oxytocin pathway, we measured oxytocin levels in plasma collected from these mice. Results showed significantly higher oxytocin levels in both S2 and S1 compared to C1 (S1, p<0.01; S2,p<0.01). Although difference was not significant, S1 also showed higher oxytocin levels compared to the control C2(Fig.1O). Furthermore, a significant mild Pearson correlation (R=0.52,*p*=2.4e-06) was observed between oxytocin levels and active contracting time (Fig.1P), highlighting a possible link between oxytocin and tameness behaviour.

### Gut microbiome repertoire in mice of selected and non-selected groups

To examine the gut microbiota of the selected and non-selected groups, shotgun metagenomic sequencing analysis was performed on 80 faecal samples collected from each mouse after the tameness test. On average, we obtained 31 million paired reads per sample. Metagenome-assembled genomes (MAGs) were used to analyse the gut microbiome composition of WHS mice.

Because the gut microbiota is a complex flora, we employed two assemblers and multiple binning approaches to obtain as many good-quality MAGs as possible (Fig.S2A). Using these approaches, we generated 14,816 MAGs (details of MAGs obtained from each binner are shown Fig.S2B, C). After removing the redundant MAGs, we obtained 374 bacterial MAGs that were more than 50% completeness and had less than 10% contamination (Fig.2, S2E, Table S3). Of these MAGs, 226 were high-quality as they were more than 90% completeness and less than 5% contamination (Fig.S2D). These 374 MAGs spanned 11 phyla, with 281 MAGs belonging to the *Bacillota A* phylum (Fig.2, Table S3). There are multiple mouse metagenome catalogues available^78,79^, with the most recent being the Mouse Gastrointestinal Bacteria Catalogue (MGBC)^80^ and the Comprehensive Mouse Microbiota Genome Catalogue (CMMGC)^70^. For our analysis, we used the CMMGC to compare our MAG data because it covers a higher number of species. Upon comparison, we identified 27 species level novel MAGs that have not been previously reported. This is further confirmed by “taxonomic check” in DFAST^72^. Of these 27 MAGs, one each belonged to *Actinomycetota*, *Bacillota B*, and *Patescibacteria*, eight to *Bacillota*, and 16 to *Bacillota A* (Fig.2). Additionally, despite the efficacy of machine learning binners in deriving higher quantity and quality MAGs, their limitations in binning all bacteria types were identified, suggesting that the application of combined methodology that integrates both single assembly and co-assembly, as well as the application of multiple binning tools is advantageous (Fig.S2E, F).

**Fig. 2.**
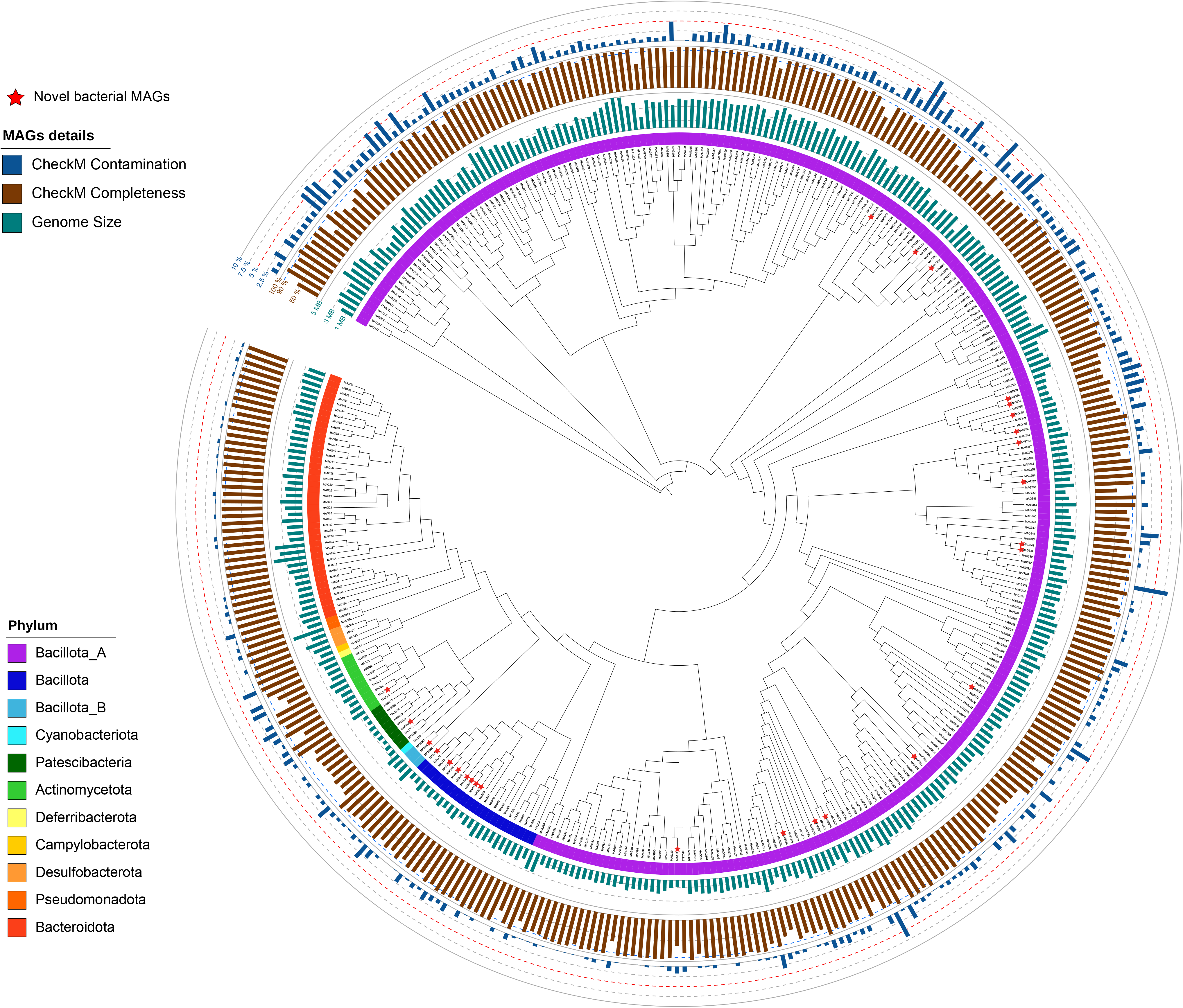
Cladogram of 374 MAGs generated. (A) All 374 MAGs were placed in this cladogram generated by GTDB output. Different phylum was represented as different colours in the innermost circle. Second circle indicates size of each MAGs; Third circle indicates CheckM completeness and fourth circle indicates CheckM contamination. All novel MAGs are indicated as red star.

For taxonomic analysis, to capture the full diversity of WHS mice gut microbiome, we used the full NCBI/RefSeq prokaryote genome sequence database (NCBI RefSeq Complete V205)^48^. We identified 665 bacterial species spanning 11 phyla, 21 classes, 45 orders, 103 families, and 339 genera (Fig.3A). Of these, 427 species were common to all four groups and contributed 99.8% abundance in all samples (Fig.3B).

**Fig. 3.**
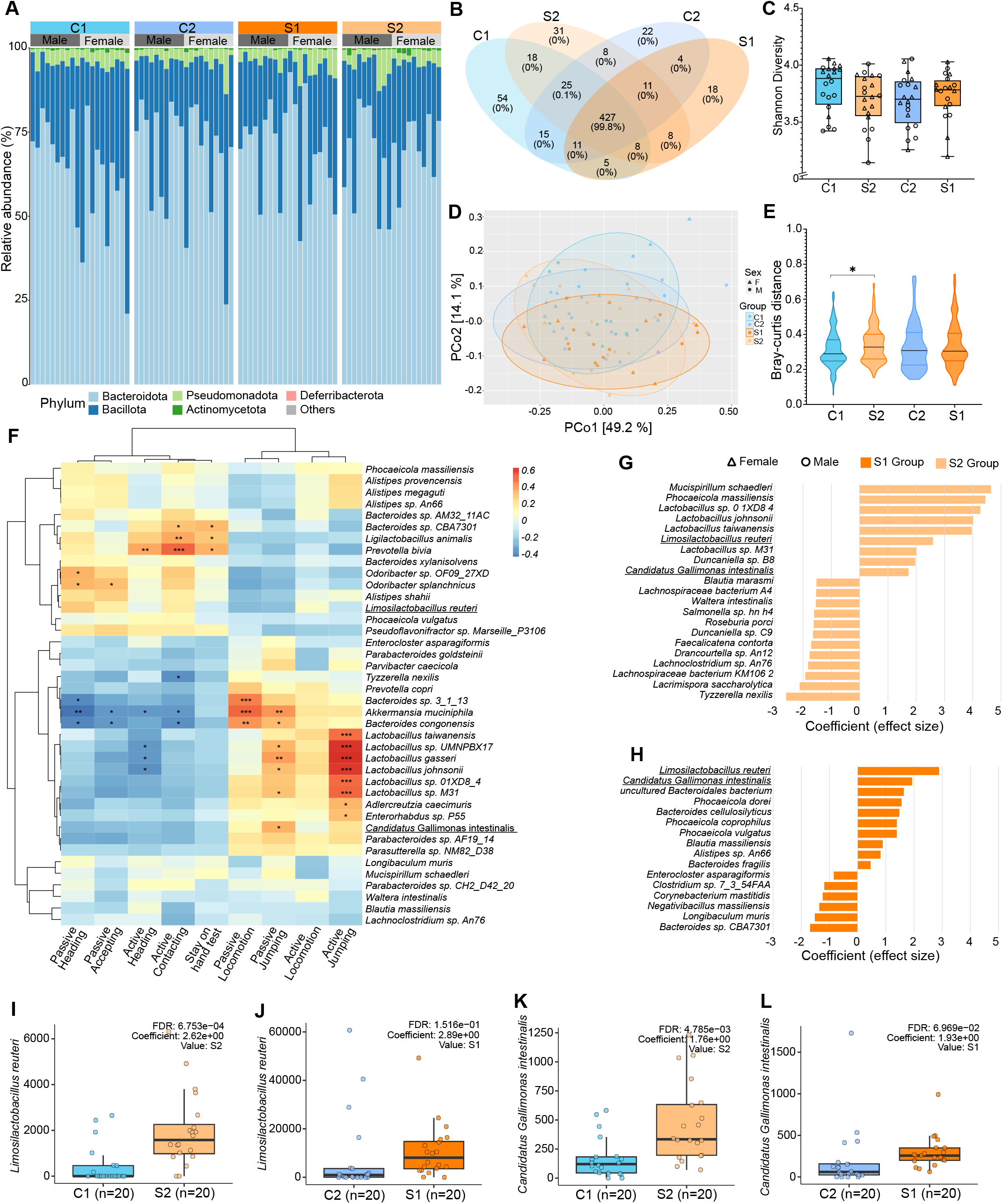
Gut microbiome diversity is similar in all groups. (A) Phylum level relative abundance in all groups, five most abundant phylum is shown; full dataset includes 11 phyla, 21 classes, 45 orders, 103 families, and 339 genera, (B) Ven diagram of 665 bacterial species present in WHS mice gut, (C) Shannon diversity, (D) Beta diversity based on Bray–Curtis dissimilarity, (E) Bray curtis distance between each groups, (F) Hierarchical clustering using Pearson correlation distance between top 40 significantly different taxa (result of random forest analysis) with score of parameters of tameness test. Significant correlation is marked with “*” (G) Association analysis in MaAsLin2 between S2 group and bacterial abundance, (H) Association analysis in MaAsLin2 between S1 group and bacterial abundance, (I-J) MaAsLin2 generated graph of *Limosilactobacillus reuteri*, (K-L) MaAsLin2 generated graph of *Candidatus* Gallimonas intestinalis. N= 80 (20 in each group with 10 male and 10 female). (*p<0.05); ***p<0.001).

In terms of alpha diversity, we observed similar Shannon diversity across all groups (Fig.3C). Taxonomic beta diversity, calculated using Bray-Curtis dissimilarity, also showed no significant differences between the groups (Fig.3D). However, when comparing the mean Bray-Curtis distance, we found a significant difference between groups C1 and S2 (p<0.01) (Fig.3E). Overall, these findings suggest that host selection pressure for tameness behaviour does not consistently affect the overall taxonomic diversity of the gut bacterial microbiota in mice.

We conducted Pearson correlation analysis on the tameness parameters scores with 40 significant taxa identified from the random forest analysis. This analysis highlighted that several *Lactobacillus* bacteria showed a significant positive correlation with both active and passive jumping behaviours, parameters related to wildness, while showing a significant negative correlation with active heading. On the other hand, *Bacteroides sp.* CBA7301, *Ligilactobacillus animalis*, and *Prevotella bivia* were significantly positively correlated with our selection pressure- active contacting (Fig.3F).

The functional analysis of our metagenomic data uncovered 410,083 gene families across all samples. Among these, 240,161 gene families were common to all four groups, indicating a core set of functionalities shared across the study subjects. Additionally, each group exhibited unique gene families, highlighting the diversity within the gut microbiota’s functional potential (Fig.S3A). Subsequently, we examined the diversity of gene families in all four groups of mice. Chao1 richness showed that the C1 group had the highest functional richness among all groups and significantly higher richness than the S1 group (p<0.01) (Fig.S3B). Unlike taxonomic diversity, the Shannon diversity of the gene family in the S1 group was significantly lower than that in the C1 group (Fig.S3C). However, beta diversity did not differ between groups (Fig.S3D). Gene pathways were also identified from gene families using the MetaCyc database.

Overall, we identified 378 gut microbial pathways in WHS mice, 263 of which were present in all four groups (Fig.S3E). Association analysis was performed on the MetaCyc pathways identified by functional analysis. The metabolic pathways in both S1 and S2 were compared with their respective controls using MaAsLin2. Overall, we identified 88 pathways in the S2 group and 49 pathways in the S1 group that met the q-value cutoff criteria. Among these, 35 pathways exhibited consistent trends in both the S1 and S2 groups, as outlined by star (*) in Table S4. Notably, we discovered several pathways related to amino acid biosynthesis and energy generation that were enriched in the selected groups. This enrichment suggests that the selection for tameness may influence metabolic functions related to these pathways, potentially contributing to the observed behavioural traits.

### The selected groups exhibited abundance in *Limosilactobacillus reuteri*

To identify the enrichment of bacterial species in tame mice and make the association more robust, we compared each selected group with its respective control group. For this analysis, we used MaAsLin2 to associate species-level gut microbiome composition with the WHS mouse group (Fig.3G, H). For MaAsLin2 analysis, only species that were significantly associated in both groups were considered to differ in abundance. We found that *Limosilactobacillus reuteri* (Fig.3I, J), previously known as *Lactobacillus reuteri*, one of the lactic acid bacteria, and *Candidatus* Gallimonas intestinalis (Fig.3K, L) were enriched in both selected groups compared to their respective controls.

Among the 374 MAGs generated, one high-quality MAG from *L. reuteri* and six MAGs of *Candidatus* Gallimonas genus were obtained. To quantify gut bacterial abundance, we used CoverM for mapping reads to all MAGs, with mapping rates ranging from 75% to 93% (Fig.S4A). Similar to Fig.3 (the Kraken2 results), we found that *L. reuteri* was significantly enriched in both selected groups compared to non-selected groups (S1, p<0.05; S2, p<0.001) (Fig.S4 B, C). However, the relative abundance of the six *Candidatus* Gallimonas MAGs did not show a significant difference between the selected and non-selected groups (Fig.S4D-I). These results combined with those of the Pearson correlation analysis performed earlier (Fig.3F) indicate that *L. reuteri* is the only bacterium that is consistently associated with tameness across our analyses. This association suggests that *L. reuteri* is linked with increased active tameness in the selected groups.

### The selected groups have a higher level of plasma pyruvate

The influence of the gut microbiota on animals occurs through the production of metabolites that are absorbed into the host and change its behaviour. To identify the metabolites that may contribute to changes in tameness, we conducted a metabolomic analysis of plasma samples obtained from WHS mice. From the capillary electrophoresis time-of-flight mass spectrometry (CE-TOFMS) measurements, 281 peaks (179 in cation mode and 102 in Anion Mode) were detected and annotated according to HMT’s standard library and the known-unknown peak library. Among the target metabolites, 70 (46 in cation mode and 24 in Anion Mode) were detected and quantified (Table S5). Correlation clustering of 70 metabolites did not show any visible difference between the control and selected groups (Fig.4A). Of these, four metabolites were significantly higher in selected mice (Fig.4B-E). To strengthen our findings, we performed MaAsLin2 analysis and after FDR correction, only pyruvic acid remained significantly associated with selected mice (TableS5). The plasma lactic acid concentration, which maintains homeostasis with pyruvic acid in the body, was also observed and found to be not significantly different between the selected and control groups (Fig.4F).

**Fig. 4.**
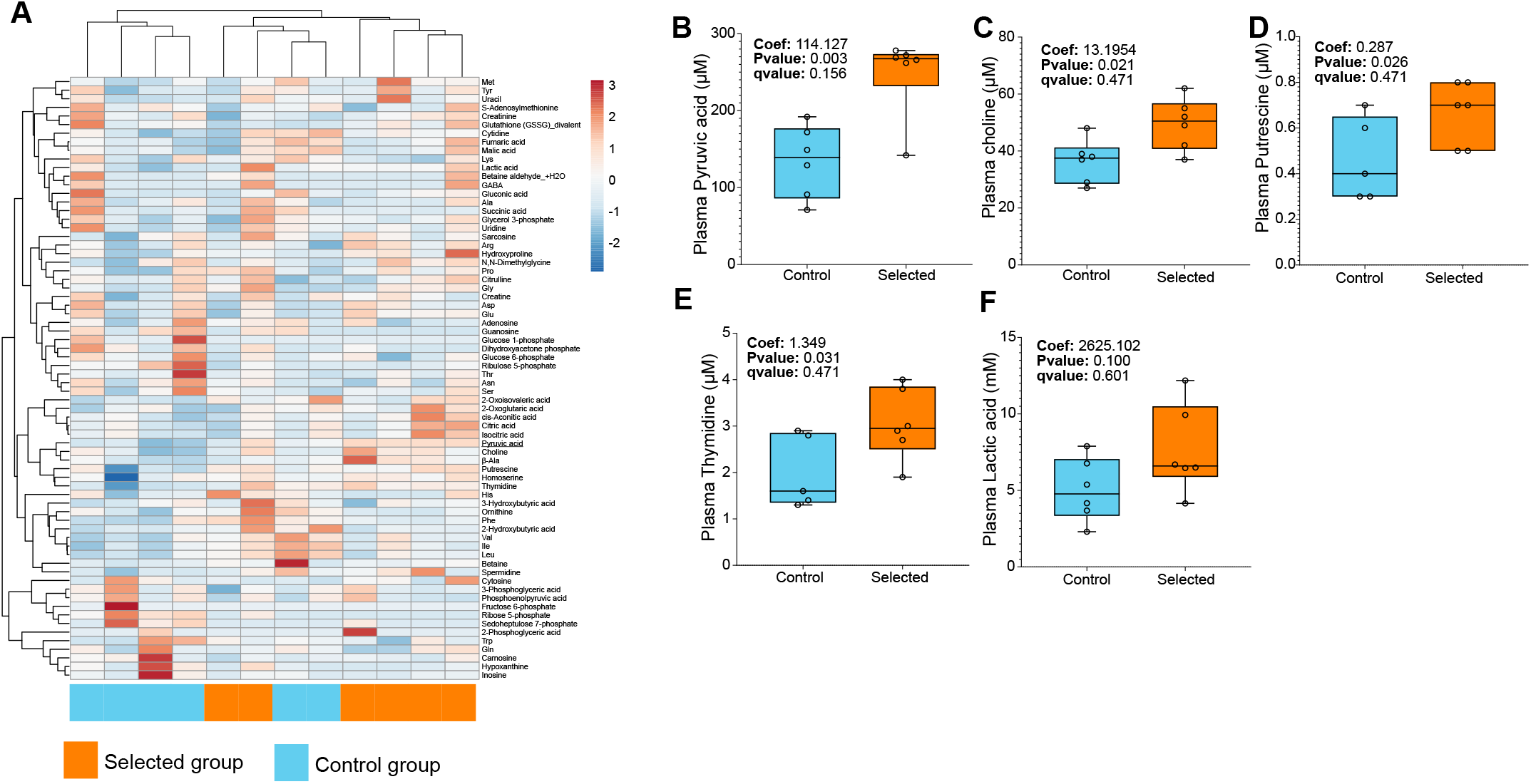
Pyruvic acid is significantly high in selected mice. (A) Heatmap with hierarchical clustering using Pearson correlation distance of 70 metabolites identified, (B) Plasma pyruvic acid, (C-E) Significantly higher metabolites before FDR correction in MaAsLin2, (C) Plasma choline, (D) Plasma putrescine, (E) Plasma thymidine, (F) Plasma lactic acid, N = 12 (6 in each group).

### *L. reuteri* administration increase tameness behaviour in non-selected mice

To examine the effects of *L. reuteri* and pyruvate in mice, strains that secrete more and less pyruvate were isolated and administered through drinking water. We isolated the bacterium from cecum material and faeces of the selected group of WHS mice. We isolated 22 *L. reuteri* strains using a specialized media with raffinose and found that 16 colonies secreted pyruvate into GAM culture media via biochemical assays (Fig.5A). *Lactobacillus helveticus* JCM1120, a pyruvate- secreting bacterial species^81^, was used as a positive control of pyruvate secreting strain. Additionally, we assessed D-lactate and L-lactate secretion due to their homeostatic relationship with pyruvate (Fig.5A). In the experiment of bacterial strain administration to C1 group mice, we chose NIG-A41 (high-pyruvate secreting strain), NIG-23 (low-pyruvate-secreting strain), as well as the *L. helveticus* JCM1120. After administration of the cultured bacterial strains for 21 days through drinking water to non-selected C1, tameness tests were conducted (Fig.5B). Mice treated with pyruvate secreting *L. reuteri*, NIG-A41, showed significantly increased active tameness compared to the PBS-administered group (p<0.05; Movies S3, S4) and *L. helveticus* administered group (p<0.05) (Fig.5C). Despite being insignificant, mice treated with less pyruvate-secreting strain, NIG-23, showed higher levels of active tameness compared to the mice treated with PBS. However, mice treated with *L. helveticus* did not show elevation in the active tameness compared with the PBS-treated mice. Active heading was also higher in NIG-A41 treated group but the difference was not significant (Fig.5D). Other behavioural parameters, stay-on-hand test, passive heading, and passive accepting did not show prominent increase in NIG-A41 treated group (Fig.5E-G). Serum pyruvate level was higher in *L. helveticus* treated group (p<0.05) but neither NIG-A41 nor NIG-23 treated mice showed any difference in serum pyruvate levels compared to the PBS treated group (Fig.5H). We also examined the colonization of these bacterial strains in host mice by quantitative PCR analysis of the bacterial DNA obtained from faeces. Significantly higher levels, more than 1500 times in average, of the *L. reuteri* genomes were found in the faeces of both NIG-A41 (p<0.01) and NIG-23 (p<0.001) treated mice compared with the PBS-treated mice (Fig.5I). In contrast, the mice treated with *L. helveticus* JCM1120, which was originally isolated from Emmental (Swiss) cheese^82^, did not show significant increase in the mice faeces (Fig.5J). We also monitored body weight changes and water intake during the administration period, but no significant differences among the groups were observed (Fig.S5A-D). Because a previous study reported that the daily administration of *L. reuteri* results in increased blood oxytocin levels^83^, we examined the oxytocin levels in these mice. Serum oxytocin levels were significantly higher in mice treated with NIG-A41 (p<0.01) compared to those in the PBS-treated group, but there was no significant difference in the mice treated with NIG-23 (Fig.5K). In addition, there is a significant mild Pearson correlation between serum oxytocin concentration and active contacting time (R=0.44, p=0.0075) (Fig. 5L). These results showed that the long-term administration of a specific strain of *L. reuteri*, NIG-A41, resulted in a higher level of active tameness with higher blood oxytocin levels, as well as higher number of faecal *L. reuteri*.

**Fig. 5.**
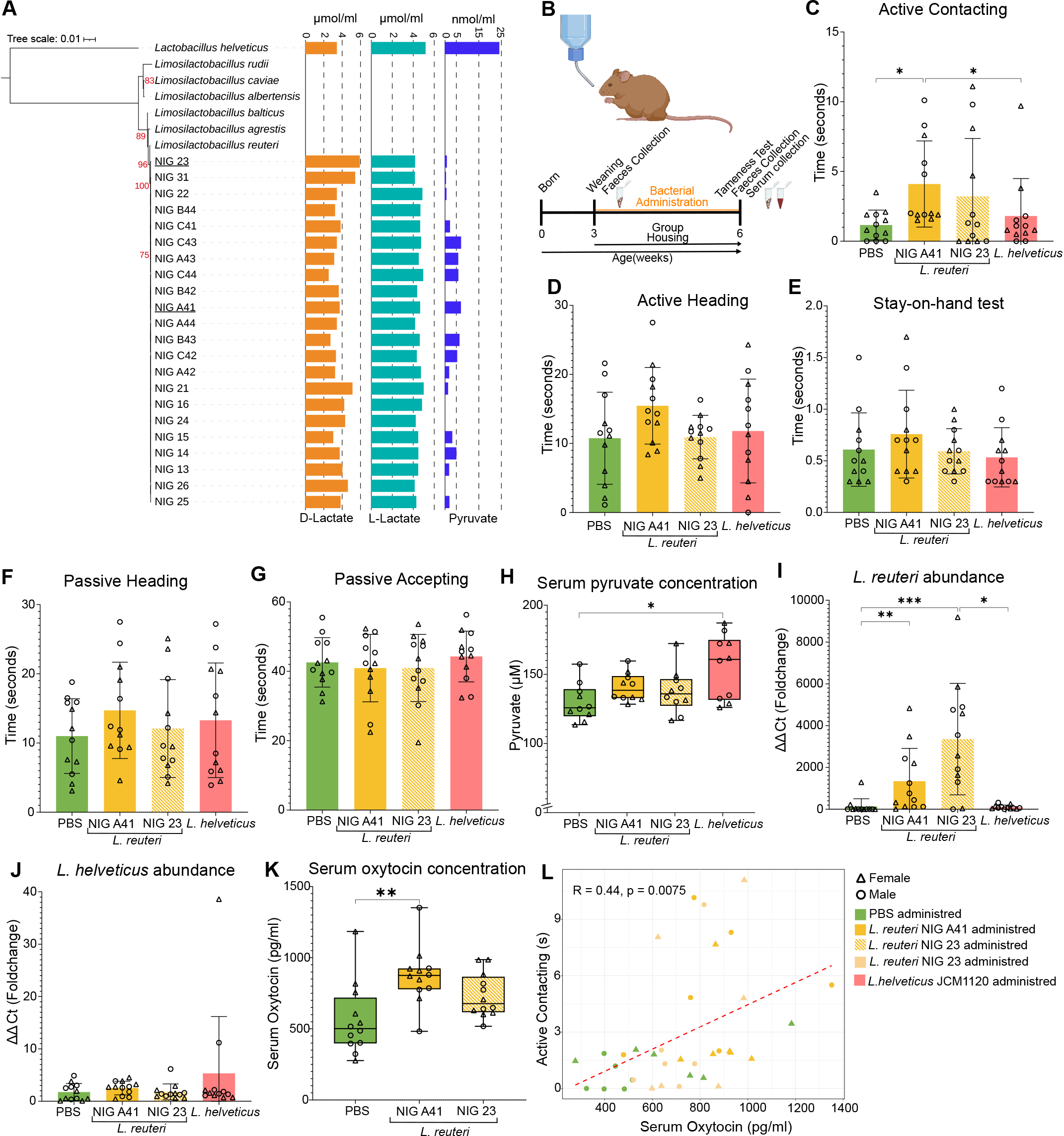
*Limosilactobacillus reuteri* administration can increase tameness behaviour in mice. (A) Maximum likelihood tree of 16S rDNA region; different species of *Limosilactobacillus* genus was used for identification and *Lactobacillus helveticus* was used as outgroup. 1000 bootstrap was run and bootstrap value of more than 70% is shown with red ink. Bar graph represent pyruvate and lactate secretion in GAM media by different colonies of *L. reuteri* isolated in current study and *L. helveticus*, (B) scheme of bacteria administration to control mice through drinking water; (C-G) Different tameness test parameters after bacterial administration, (C) Active contacting, (D) Active heading, (E) Stay on hand test, (F) Passive heading, (G) passive accepting, (H) Serum pyruvate level after bacterial administration, (I) qRT-PCR quantification of *L. reuteri* present in faeces; (J) qRT-PCR quantification of *L. helveticus* present in faeces; (K) Serum oxytocin concentration, (L) Pearson correlation between oxytocin concentration and active contacting time. N= 48 (12 in each group with 6 male and 6 female) in C-G, I, and J, N=40 (10 in each group with 5 male and 5 female) in H, and N= 36 (12 in each group with 6 male and 6 female) in K and L. (*p<0.05; **p<0.01; ***p<0.001). Bar graphs show means ± SD with individual data points.

## Discussion

In this study, we investigated the relationship between gut microbiota and active tameness behaviour and investigated its potential underlying mechanisms. We studied the gut microbiota in selected groups of mice that exhibited high tameness and in the non-selected groups of mice that exhibited low tameness. Our findings reveal that while selection for tameness does not markedly change the taxonomic or functional diversity of the gut microbiota, but leads to the enrichment of one specific bacterial species, *L. reuteri*, in tame mice. Furthermore, our plasma metabolic analysis revealed a significant elevation of pyruvate levels in tame mice. The administration of pyruvate- secreting *L. reuteri* increases blood oxytocin levels and active tameness in non-selected mice. We did not observe a consistent difference in gut bacteriome alpha or beta diversity between control and selected mice, suggesting that selective breeding for tameness does not influence these diversity metrics (Fig.3). However, a separate analysis focusing on virome sequences from the same dataset showed a significant difference in beta diversity between control and selected mice^84^. At the species level, several studies have associated bacterial genera and species with domestication traits in animals. Laboratory mice show higher levels of

*Akkermansiaceae*, *Streptococcaceae*, and *Enterobacteriaceae* than wild types^11^. Domesticated horses exhibit more archaea than feral ones^13^, while domesticated buffalo have enriched Cyanobacteria and TM7 phyla^14^. In chickens, low fear selection increases *Clostridiales* and *Bacteroidales*, while high fear boosts *Lactobacillales* population^16^. These reports suggested that a factor other than host genetics, the gut microbiota, is associated with the domestication process. Although fully isolating experiments from external influences is challenging, uniform conditions for WHS mice helped mitigate this issue, revealing a link between tameness selection and *L. reuteri* abundance (Fig.3).

The gut microbiota can alter host energy and lipid metabolism by producing energy metabolites, such as pyruvate, fumaric acid, and citric acid, and by influencing triglyceride levels in the host plasma^85^. Pyruvate originating from the gut microbiome is absorbed by the host and contributes to the priming of the immune system and protection against *Salmonella* infection. This pyruvate is secreted by *L. helveticus*^81^, a species which belongs to the same family (*Lactobacillaceae*) as *L. reuteri*. High circulating pyruvate levels in the blood and brain may also protect against Alzheimer’s disease^86^ by reducing age-related cognitive decline^87^. In the present study, we found that mice with high active tameness had high plasma pyruvate levels (Fig.4B).

Twenty-two strains of *L. reuteri* were isolated by culturing colonies on selective media, sixteen of which showed pyruvate secretion into GAM medium, which is a novel finding^81^. Previous studies have linked *L. reuteri* administration to altered social behaviours in mice^88–90^. Our study showed that pyruvate-secreting *L. reuteri* increased active tameness, although pyruvate was not interpreted as a direct cause. We also found that bacteria isolated from the same host species exhibited more optimized colonization, without the sex-based colonization differences reported by Donovan et al.^90^. Administration of *L. reuteri* through drinking water surged the populations over 1500-fold (Fig.5I), and this large change of *L. reuteri* levels likely contributed to the observed increase in tameness.

Our results underscore the host genetics’ pivotal role in tameness, with C1 mice showing a less pronounced tameness level after *L. reuteri* administration (Fig.5C) compared to S1 and S2 groups (Fig.1G). This indicates that while the gut microbiota may influence behaviour, changes in host genetic factors are a major factor for changes in tameness during the selective breeding. Despite these facts, the result that long-term administration of live *L. reuteri* via drinking water can significantly increase the behaviour of tameness by vastly increasing its population in the gut is noteworthy for future studies.

The role of *L. reuteri* in influencing tameness behaviour warrants further investigation to elucidate the underlying mechanisms. Previous research by Matsumoto et al.^21^ indicated that in WHS mice selected for tameness, there is higher expression of the gene for the oxytocin receptor— a receptor for the social bonding neuropeptide hormone, oxytocin, in the hippocampus. In addition, tameness selection was found to lead to increases in sociability in WHS mice^77^. Similarly, a previous study observed an increase in plasma oxytocin levels in C57BL/6 mice treated with *L. reuteri* through their drinking water^83^. The administration of *L. reuteri* was also associated with an increase in oxytocin-positive neurons in the paraventricular nucleus of the hypothalamus^88,89^. Sgritta et al.^91^ demonstrated that *L. reuteri* communicates with the brain, thereby affecting social behaviour via the vagus nerve. Buffington et al.^92^ identified that the gut microbiota metabolite tetrahydrobiopterin, enhanced by *L. reuteri*, can alleviate social deficits in germ-free mice, a process linked to increased oxytocin release^93^. Danhof et al. ^94^ discovered that oxytocin is produced and secreted by intestinal epithelium enterocytes, triggered by *L. reuteri*. These collective findings pave the way for a hypothesis that oxytocin might be a central player in how *L. reuteri* influences tameness behaviour. In our study, basal oxytocin levels were higher in the selected groups compared to the non-selected groups, and *L. reuteri* administration boosted serum oxytocin in the non-selected (C1) mice (Fig.5K). A previous study showed that oxytocin also increases pyruvate dehydrogenase activity in adipocytes^95^ and enhances gluconeogenesis in hepatocytes^96^, both of which could lower pyruvate levels. This may explain why no increase in serum pyruvate was detected in *L. reuteri*-treated mice in the current study (Fig.5H).

Overall, our findings contribute to the growing body of knowledge about the complex relationships between behaviour, the host genome, and the gut microbiota in the context of animal domestication. The potential link between *L. reuteri* and oxytocin pathways opens new avenues for exploring the biological basis of domestication and tameness in animals.

## Acknowledgements

We appreciate, Motoko Nihei, Akiko Tsuchiya, Kyouhei Kurosawa, and Kazumichi Fujiwara for their technical assistance. Computations were partially performed on the NIG supercomputer at ROIS National Institute of Genetics. We would like to thank Editage (www.editage.jp) for English language editing.

## Authors’ contributions

B.B.B. and T. K. conceived the project and designed the experiments; B.B.B. performed the experiments and analysed the data; A.T. performed shotgun sequencing; H.M. and K.K. provided intellectual insights and guidance. T.K. provided supervision and funding; The initial draft of the manuscript was authored by B.B.B. and T.K., with the final Ver. being written collaboratively, incorporating feedback from all authors. Each author has approved the final version of the manuscript.

## Availability of data

All data required to evaluate the conclusions of this study are presented in the paper or supplementary materials. All 80 raw shotgun metagenomic sequencing datasets generated in this study are available in the NCBI under BioProject PRJDB15857 with BioSample accession numbers from SAMD00614304 to SAMD00614383 and SRA accession numbers from DRR480456 to DRR480535. All 16S rDNA sequences generated in this study can be accessed using accession IDs from LC801559 to LC801580 (Please use this link to search; https://www.ncbi.nlm.nih.gov/nuccore/). Non-redundant set of MAGs (374) including novel ones are available in zenodo database (https://doi.org/10.5281/zenodo.8289507). This paper does not report original code. Code used in this study are available at GitHub repository (https://github.com/bhimbbiswa/Gut-microbiota-influence-on-animal-domestication).

