## Supplementary Figures for "Enhanced tameness by *Limosilactobacillus reuteri* from gut microbiota of selectively bred mice"

### Supplementary Materials

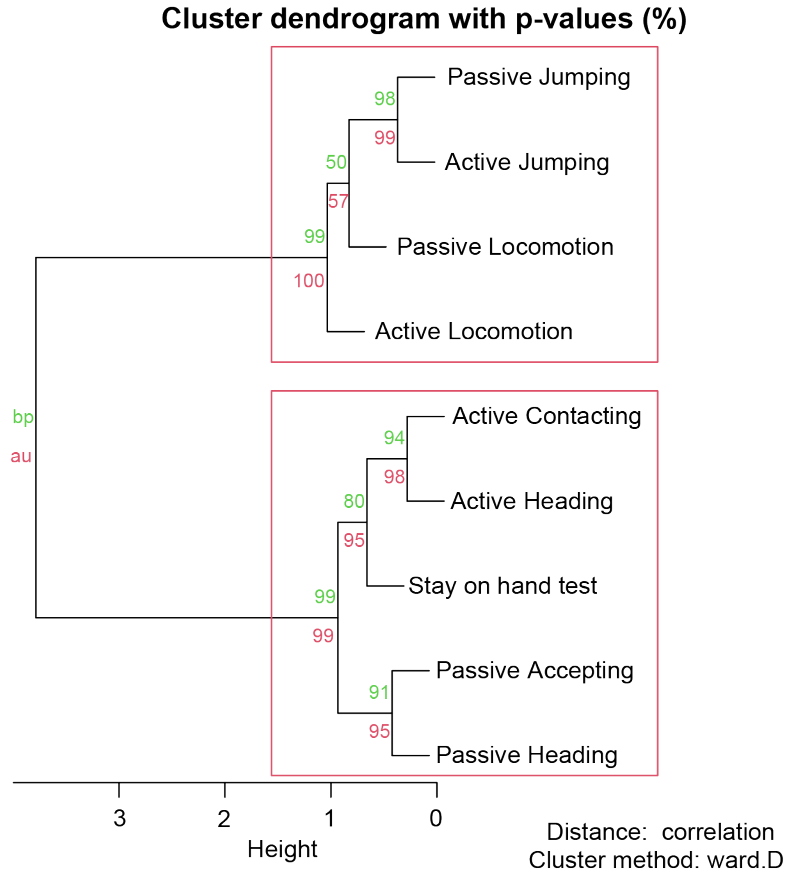


**Figure S1: Correlation clustering of tameness test parameters.** All nine parameters were normalized and clustered using the correlation distance and Ward's method. The red rectangle represents significant clusters based on the p-value calculated using pvclust. 'au' stands for (Approximately Unbiased) p-value and 'BP' for Bootstrap Probability values.

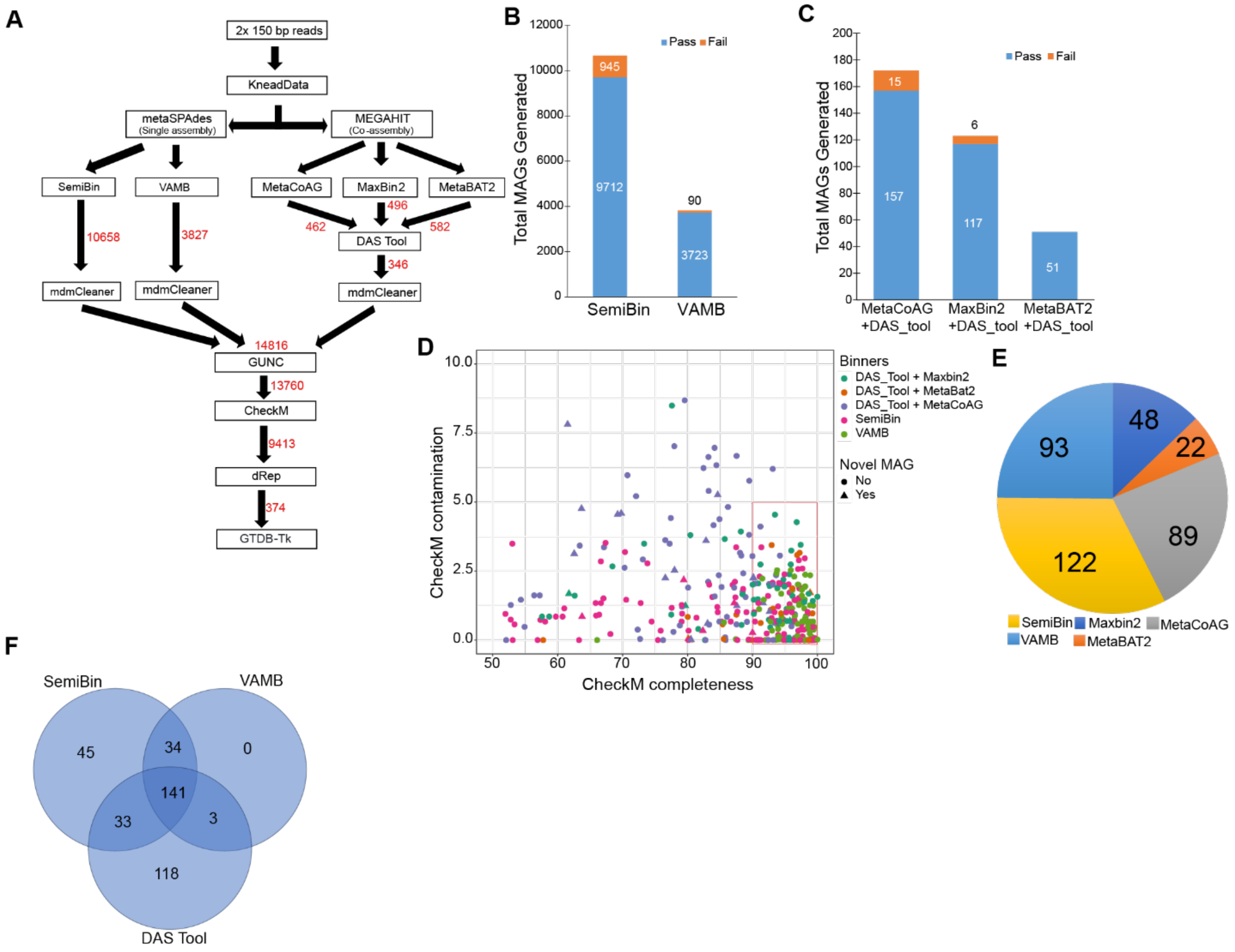


**Figure S2: Details about MAGs generated (A)** Scheme for high quality metagenome assembled genome (MAGs) generation, MAGs generated were shown next to each step with red ink, **(B-C)**Total MAGs generated by each binner with information about number of MAGs passing GUNC chimeric test, **(D)** CheckM result of all 374 MAGs obtained, red rectangle encompasses MAGS which have >90% completeness and <5% contamination, **(E)** Distribution of origin of 374 MAGs generated in the current study, **(F)** Ven diagram of origin of MAGs from same cluster of dRep (These data were compiled from clustering result of dRep where a MAG is considered to be of the same species if they cluster together at 95% ANI).

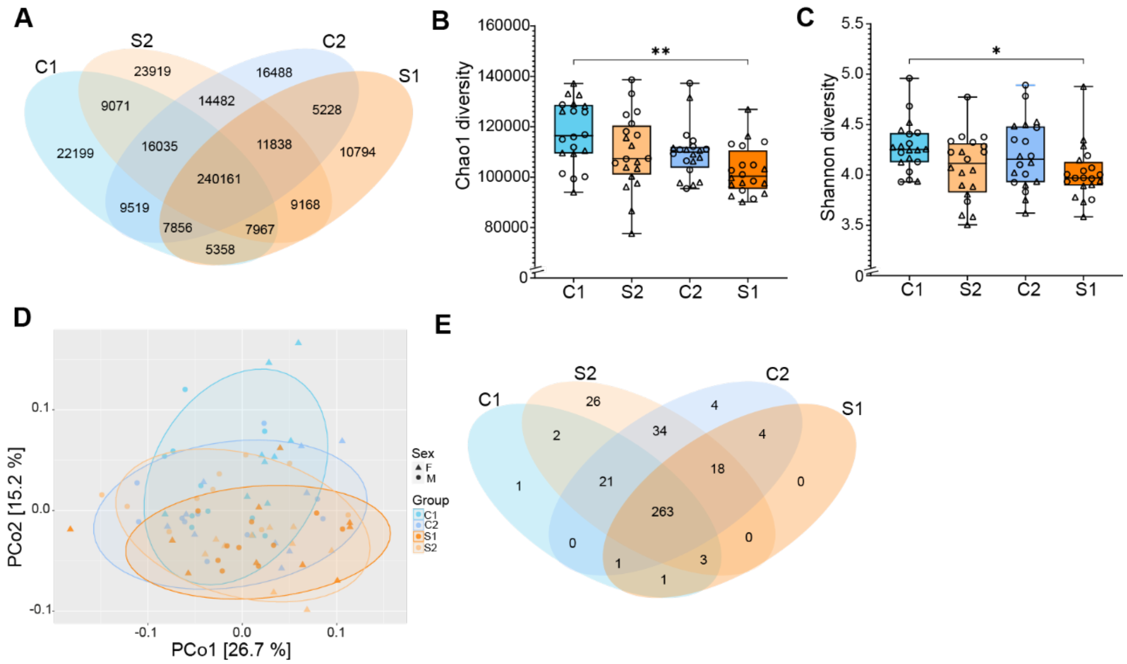


**Figure S3: Host tameness selection pressure does not change gut microbiota functional diversity (A)** Venn diagram showing all gene family identified; **(B)** gene family Chao1 diversity, **(C)** Gene family Shannon diversity; **(D)** Beta diversity based on Bray–Curtis dissimilarity. **(E)** Venn diagram of gut microbiota metabolic pathways identified. N= 80 (20 in each group with 10 male and 10 female). (*p<0.05); **p<0.01; ***p<0.001).

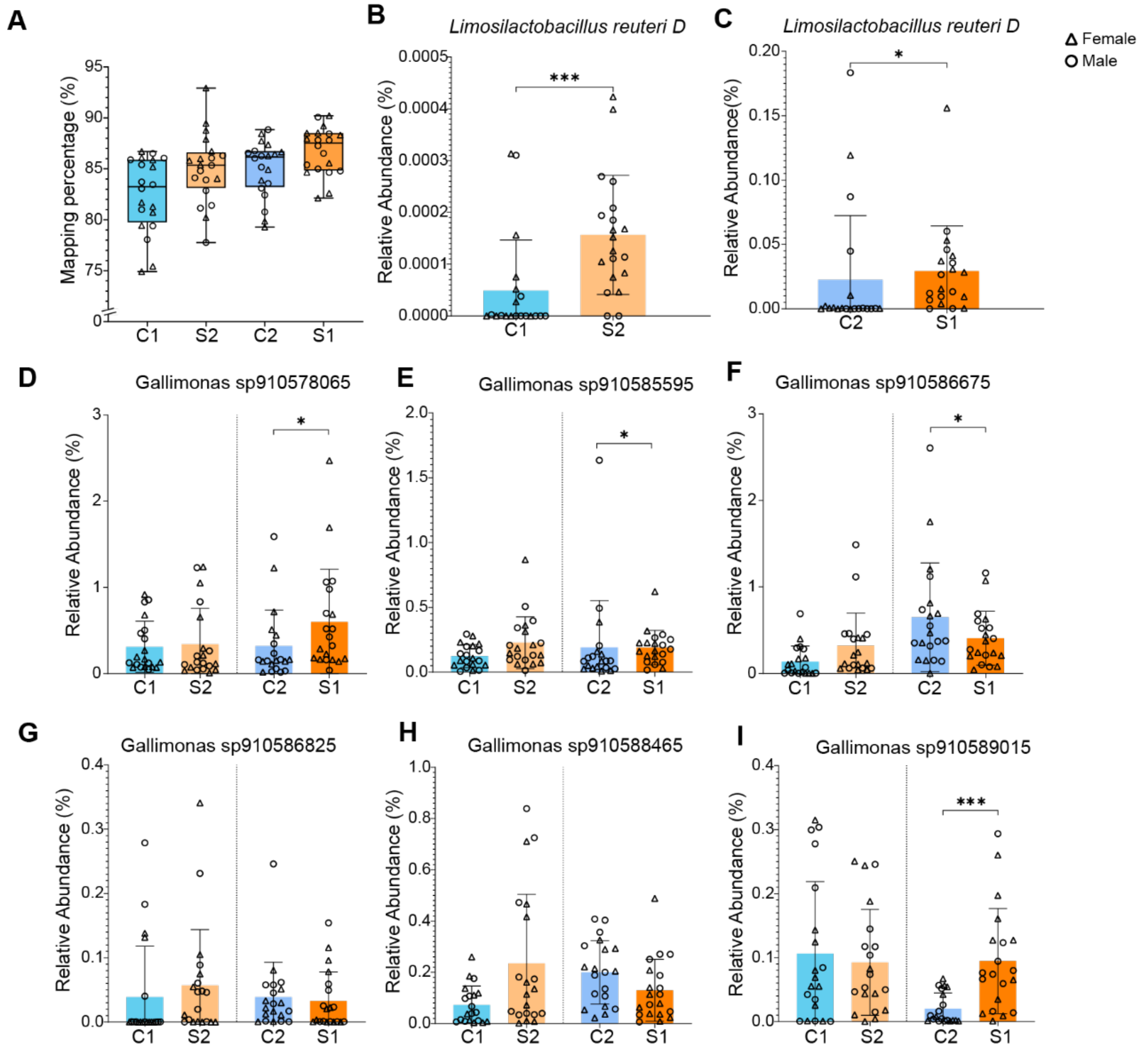


**Figure S4: Relative abundance of MAGs** (A) Mapping percentage of each sample filtered sequences against MAGs generated in this study; (B) relative abundance of *Limosilactobacillus reuteri* MAGs in C1 & S2 (One sample was not shown as it exceeds y-axis, but utilized for statistical calculation); (C) relative abundance of Limosilactobacillus reuteri MAGs in C2 & S1; (D-I) relative abundance of six *Candidatus* Gallimonas MAGs; N= 80 (20 in each group with 10 male and 10 female). (*p<0.05); **p<0.01; ***p<0.001). Bar graphs show means ± SD with individual data points.

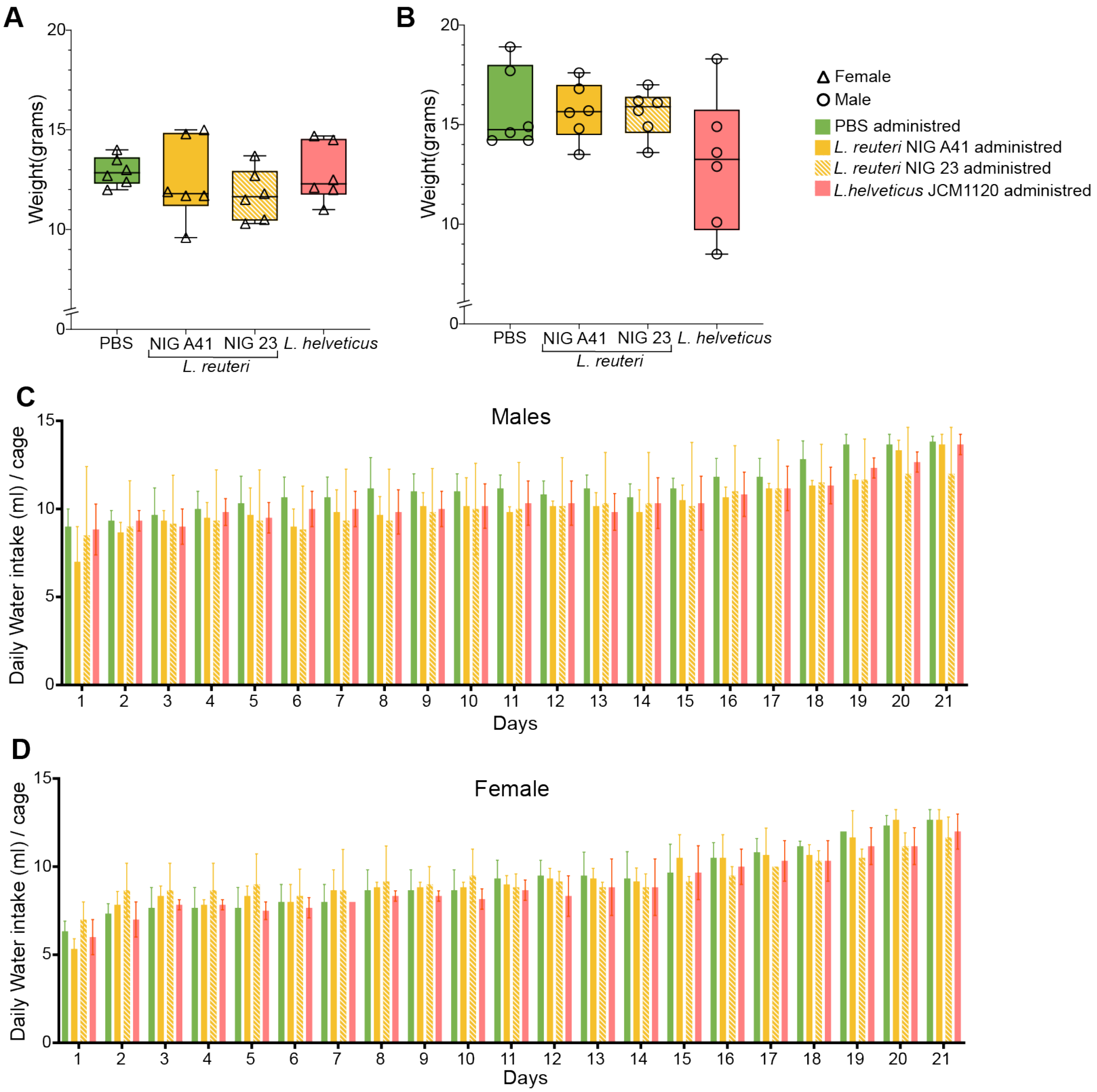


**Figure S5: Figure S4: No change in body weight and water intake due to bacterial administration, (A)** Body weight of females at 6 weeks of age after bacterial administration **(B)** Body weight of males at 6 weeks of age after bacterial administration; **(C)** Daily water intake in males for 21 days; **(D)** Daily water intake in female for 21 days. N= 48 (12 per groups). As mice were kept as same sex pairs during this experiment, so water intake data is from 6 cages per group (3 from males and 3 from females). Bar graphs show means ± SD with individual data points.
